# Non-mutated human tau stimulates Alzheimer’s disease-relevant neurodegeneration in a microglia-dependent manner

**DOI:** 10.1101/2025.01.10.632400

**Authors:** Ethan R. Roy, Qiang Wang, Kexin Huang, Sanming Li, Yuanyuan Fan, Estrella Escobar, Shuning Huang, Juan J. Herrera, Wenbo Li, Clare Pridans, Xiaobo Zhou, Cynthia Ju, Wei Cao

## Abstract

The accumulation of abnormal, non-mutated tau protein is a key pathological hallmark of Alzheimer’s disease (AD). Despite its strong association with disease progression, the mechanisms by which tau drives neurodegeneration in the brain remain poorly understood. Here, we selectively expressed non-mutated or mutated human microtubule-associated protein tau (*hMAPT*) in neurons across the brain and observed neurodegeneration in the hippocampus, especially associated with non-mutated human tau. Single-nuclei RNA sequencing confirmed a selective loss of hippocampal excitatory neurons by the wild-type tau and revealed the upregulation of neurodegeneration-related pathways in the affected populations. The accumulation of phosphorylated tau was accompanied by cellular stress in neurons and reactive gliosis in multiple brain regions. Notably, the lifelong absence of microglia significantly and differentially influenced the extent of neurodegeneration in the hippocampus and thalamus. Therefore, our study established an AD-relevant tauopathy mouse model, elucidated both neuron-intrinsic and neuron-extrinsic responses, and highlighted critical and complex roles of microglia in modulating tau-driven neurodegeneration.

## Introduction

Alzheimer’s disease (AD) is the most prevalent form of neurodegeneration with ominous societal impacts. The AD brain accumulates neurofibrillary tangles (NFTs) comprised of aggregated tau protein (encoded by *MAPT* gene), a core pathology centrally associated with the clinical manifestation (Chang et al., 2021; Wang and Mandelkow, 2016). Among “tauopathies”, neurodegenerative disorders that share excessive tau aggregation, primary tauopathies are caused by the genetic predisposition of various *MAPT* mutations (Goedert and Jakes, 2005). A secondary tauopathy, AD brains harbor abundant amyloid plaques and build up 3-8 times more tau protein than non-diseased controls, leading to the development of NFTs containing non-mutated tau (Hu et al., 2002; Khatoon et al., 1992; Knopman et al., 2021; Scheltens et al., 2021; Yamamori et al., 2007).

Abundantly expressed in neurons, tau primarily associates with and stabilizes microtubules and regulates various key neuronal processes (Chang et al., 2021; Wang and Mandelkow, 2016). To investigate AD-relevant tau-dependent neurodegeneration, we recently developed a cellular tauopathy model by overexpressing human tau (hTau) in mouse primary neurons (Li et al., 2024). This system allowed us to comprehensively assess neuronal-intrinsic responses to wild-type hTau and enabled a direct comparison with frontotemporal lobar degeneration (FTLD)-associated mutant form. Remarkably, we observed heightened neurotoxicity triggered by wild-type hTau and revealed joined MAPK-DLK signaling and DNA damage response in the degenerative process pertinent to AD (Li et al., 2024). Given the cellular complexity of the central nervous system (CNS), it is imperative to recapitulate the neurotoxic phenotype associated with non-mutated hTau *in vivo*. In the brain, glial cells, especially microglia, have been critically implicated in the pathogenesis of tauopathy as a result of mutant *MAPT* transgene overexpression (Johnson and Lukens, 2023; Parra Bravo et al., 2024; Shi et al., 2017). At this time, the contribution of non-neuronal cells or factors to AD-relevant neurodegeneration instigated by wild-type hTau is unknown.

In this study, we established a murine tauopathy with the AAV vectors used in our *in vitro* study to express human wild-type or P301L tau selectively in neurons *in vivo*. We systemically characterized the histological, structural, cellular, and molecular changes that occurred in the brains expressing these hTau forms. Furthermore, we assessed the functional requirement of microglia in wild-type hTau-induced neurodegeneration by applying this tauopathy model to *Csf1r^FIRE^* mice, a transgenic mouse strain devoid of microglia (Rojo et al., 2019). This investigation confirmed a greater neurotoxic effect of wild-type over P301L hTau to the brain and demonstrated an essential role of microglia in the progression of AD-relevant tau-mediated neurodegeneration.

## Results

### AAV-mediated human tau expression in the mouse brain

To establish a model of tau pathology *in vivo*, we inoculated neonatal C57BL/6J mice with AAV vectors containing different forms of human *MAPT*: one with an early STOP codon preventing translation of tau protein (AAV-STOP), full-length wild-type 2N4R form (AAV-Tau), and FTLD-associated P301L 2N4R mutant (AAV-P301L Tau) (Fig. 1A). At multiple timepoints, positive expression of transgenic hTau was detected throughout the hippocampal formation as well as in some surrounding regions, such as the cortex and subcortical regions like the thalamus (Fig. 1B). Driven by the human *SYN1* promoter (Li et al., 2024), hTau expression was confined to neurons (Fig. 1C). Given the moderate dose of AAV inoculated, between 10-20% of hippocampal neurons expressed hTau, whereas cortical neurons did so at a lower percentage (Fig. 1C). Within neurons, hTau was localized mostly to the soma, with some expression in dendrites and axons as well (Fig. 1C).

**Figure 1.**
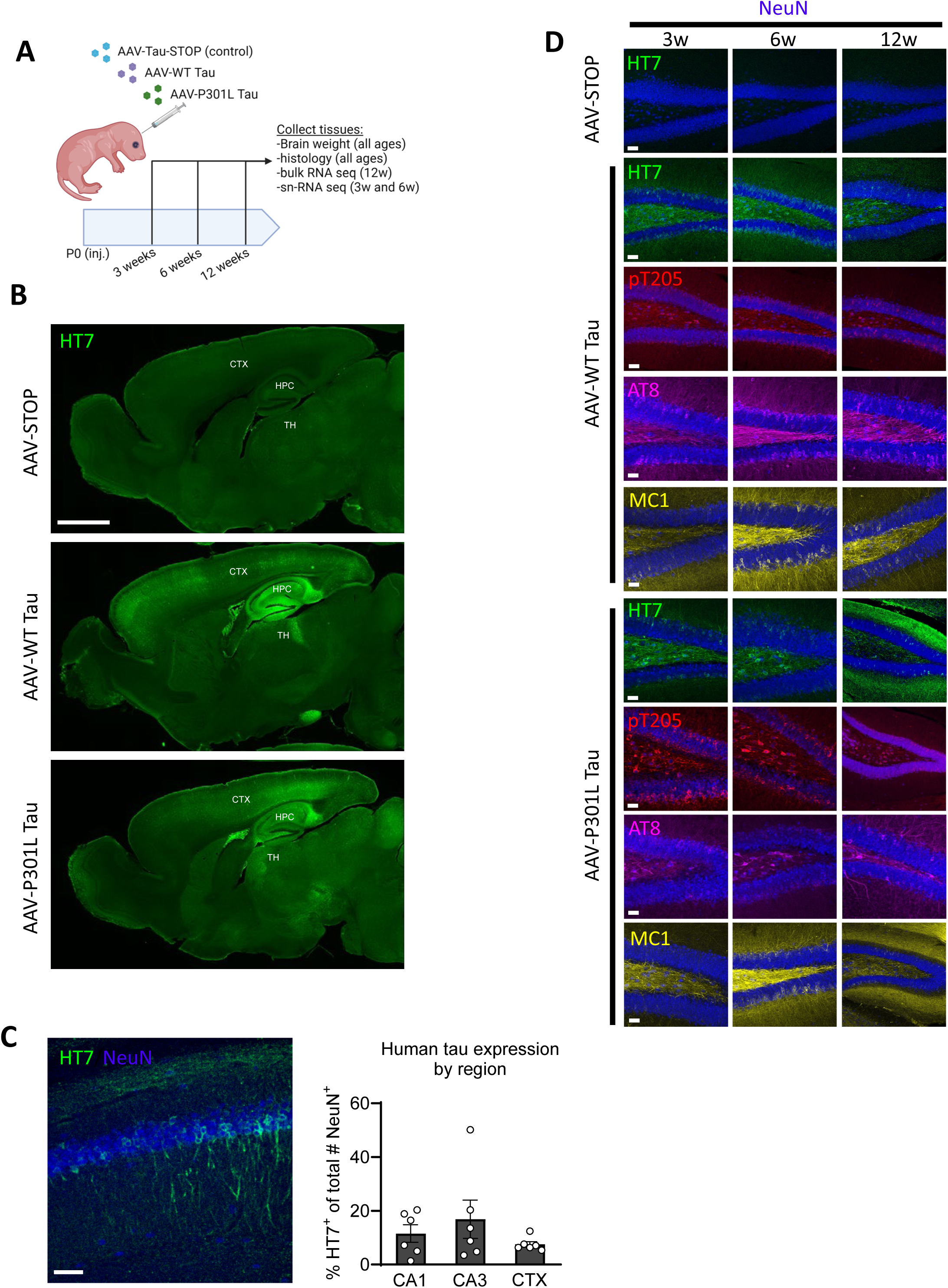
A) Schematic showing AAV-mediated overexpression of transgenic human *MAPT* forms in neurons *in vivo*. B) Fluorescence microscopy images of brain sections from animals (6 weeks of age) overexpressing transgenic human tau (HT7), persisting into adulthood. Scale bar, 1.5 mm. CTX, cortex; HPC, hippocampus; TH, thalamus. C) Representative confocal image of HT7^+^ total transgenic human tau expression in the CA1 of B6 mice (scale bar, 50 µm), with quantification of frequency of hTau^+^ neurons in multiple brain regions. D) Panel of antibodies targeting tau forms, including total transgenic tau (HT7), phospho-tau (pT205, AT8), and a conformational epitope (MC1), expressed by either wild-type or P301L mutant tau constructs. Representative images were taken from the dentate gyrus of animals from different time points. Scale bars, 50 µm.

Human tau protein undergoes post-translational modifications at various sites to aggregate and form toxic species while eventually assembling into NFTs (Dujardin et al., 2020; Goedert and Jakes, 2005; Guo et al., 2017; Tracy and Gan, 2018; Wesseling et al., 2020). To gain understanding of pathological tau formation, we assessed multiple histological markers of hyperphosphorylated tau (p-tau), including pT205, AT8 (which detects both p-S202 and p-T205), and MC1 (which detects a conformational epitope within amino acids 312-322). Robust expression of these markers was detected in multiple subregions of the hippocampal formation, especially in the dentate gyrus (Fig. 1D), the CA1, and the CA3, with both wild-type and P301L mutant tau, and at all the time points assessed. Taken together, these results demonstrate a robust, persistent, and pathological tau expression from both wild-type and mutated *MAPT* sequences in multiple brain regions.

### Brain pathologies induced by human tau

To assess how wild-type and mutated tau affect the mouse brain on the macroscopic scale, we first gathered brain weight data on a large cohort of animals harvested at different time points. While no significant difference was noticed at 3 weeks, there was a significant decrease in brain weight (∼6.7%) brought by wild-type AAV-Tau at 6 weeks (Fig. 2A). By contrast, animals expressing P301L tau did not show brain weight changes at 3 weeks, and only a very small (∼3.1%) decrease at 6 weeks (Fig. S1A), suggesting a less pathogenic impact by mutant tau protein despite comparable expression levels, consistent with our findings *in vitro* (Li et al., 2024). Of note, wild-type hTau expression in CNS had no impact on the body weight of the animals (Fig. S1B).

**Figure 2.**
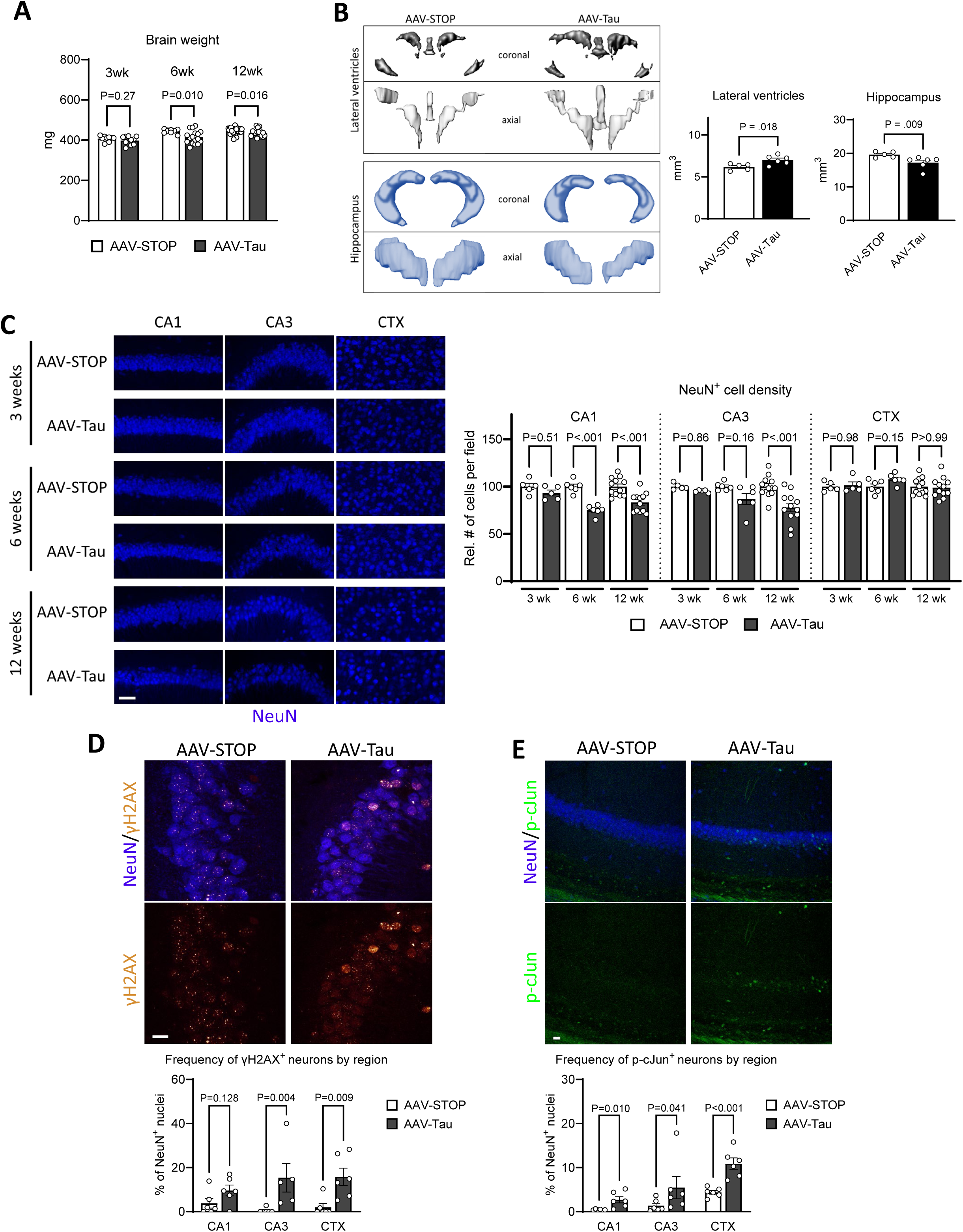
A) Quantification of brain weight at various time points in animals overexpressing wild-type tau (“AAV-Tau”) compared to non-coding control vector (“AAV-STOP”). B) 3D volumetric renderings of lateral ventricles (top) and hippocampi (bottom) from AAV-Tau and AAV-STOP animals at 3 months of age based on MRI images. Quantification of these volumes is shown. C) Representative confocal images of NeuN^+^ neurons in three brain subregions, at three time points after neonatal inoculation. Scale bar, 50 µm. Quantification of relative neuronal density in AAV-Tau compared to AAV-STOP animals by region and age. D) Representative confocal images of DNA damage in hippocampal neurons (CA3) marked by nuclear γH2AX^+^ foci. Quantification of γH2AX^+^ neurons in brain subregions at 6 weeks of age. E) Representative confocal images of DLK stress pathway in hippocampal neurons (CA3) marked by nuclear p-cJun signal. Quantification of p-cJun^+^ neurons in brain subregions at 6 weeks of age.

Because non-mutated tau has direct biological relevance to AD, we decided to focus primarily on wild-type tau (referred to throughout as “Tau”) for the remainder of this study. To examine the overall changes in brain structure brought by Tau, we performed a non-biased analysis via resting-state volumetric MRI on a cohort of mice 12 weeks after AAV inoculation. 3D reconstruction of structural volumes revealed a significant increase (∼13.4%) in total lateral ventricle volume and a concomitant decrease (∼12.2%) in total hippocampal volume (Fig. 2B). Ventricular enlargement is a prominent biomarker of AD progression and most neurodegenerative disorders (Apostolova et al., 2012). The hippocampus represents the utmost damaged brain region in AD, and its dysfunction underlies the core feature of cognitive impairment (Serrano-Pozo et al., 2011). Therefore, these findings signify a gross degenerative effect of wild-type human tau on the murine brain that is highly relevant to AD.

We next examined cellular and molecular changes in tau-expressing animals at the histological level. Using NeuN as a pan-neuronal marker, we measured the density of neurons in the hippocampus, specifically in CA1 and CA3 areas, and other brain regions, including layer V of the cortex. While the cortex did not show a tau-dependent decrease in neuronal density at any age, the CA1 and CA3 displayed progressive Tau-dependent neuron loss beginning at 6 weeks (Fig. 2C). This observation is consistent with the abundant hTau and p-tau expression in the hippocampus region and in line with the structural changes observed with brain weight and volumetric MRI.

We previously identified a functional synergy between DNA damage response and the DLK-MAPK pathway in Tau-induced neurotoxicity on cultured primary neurons (Li et al., 2024). Phosphorylated histone variant H2AX (γH2AX) is a core molecular marker of double-strand DNA damage and repair (Mah et al., 2010). Staining for γH2AX revealed neuronal nuclear foci, indicating genomic DNA damage, at a higher frequency in AAV-Tau animals compared to control in both the CA3 and cortex (Fig. 2D). Phosphorylated by c-Jun N-terminal kinases (JNKs), c-Jun protein acts as a potent transcriptional factor downstream of MAP kinase signaling (Bohmann et al., 1987). In a similar manner, CA1, CA3, and cortical neurons in AAV-Tau animals expressed a higher frequency of nuclear p-cJun, indicating activation of the DLK-MAPK injury signaling (Fig. 2E). Interestingly, these pathways were elevated at 6 weeks, but not at 3 weeks (Fig. S1C), suggesting a delayed molecular response to early Tau expression in the brain. Taken together, these results demonstrate the toxic effect of wild-type human tau in mouse brains, which potently infringes on brain structure, reduces neuronal density in the hippocampus, and activates intrinsic pathways involved in genomic stress and injury response.

### Molecular alterations in hippocampal neurons brought on by non-mutated hTau

To investigate specific molecular changes in neuronal populations during pathology induced by Tau, we performed single-nucleus RNA sequencing (snRNA-seq) on pooled hippocampi from animals receiving AAV-STOP or AAV-Tau at 6 weeks of age (n=3 per group). A total of 39,861 cells were sequenced. After preprocessing steps including quality control and normalization (see Methods), 39,568 cells with 22,374 genes were kept for the downstream analysis. Clustering with Seurat, we detected 15 distinct cell clusters, visualized by Uniform Manifold Approximation and Projection (UMAP) (Fig. S2A). Using established marker genes (Koutsodendris et al., 2023; Wang et al., 2024), six major cell types were identified in the tissue, the most prominent among them by cell counts being neurons. Other cell types in the dataset included oligodendrocytes, oligodendrocyte precursor cells (OPCs), astrocytes, microglia, and vascular cells (Fig. 3A).

**Figure 3.**
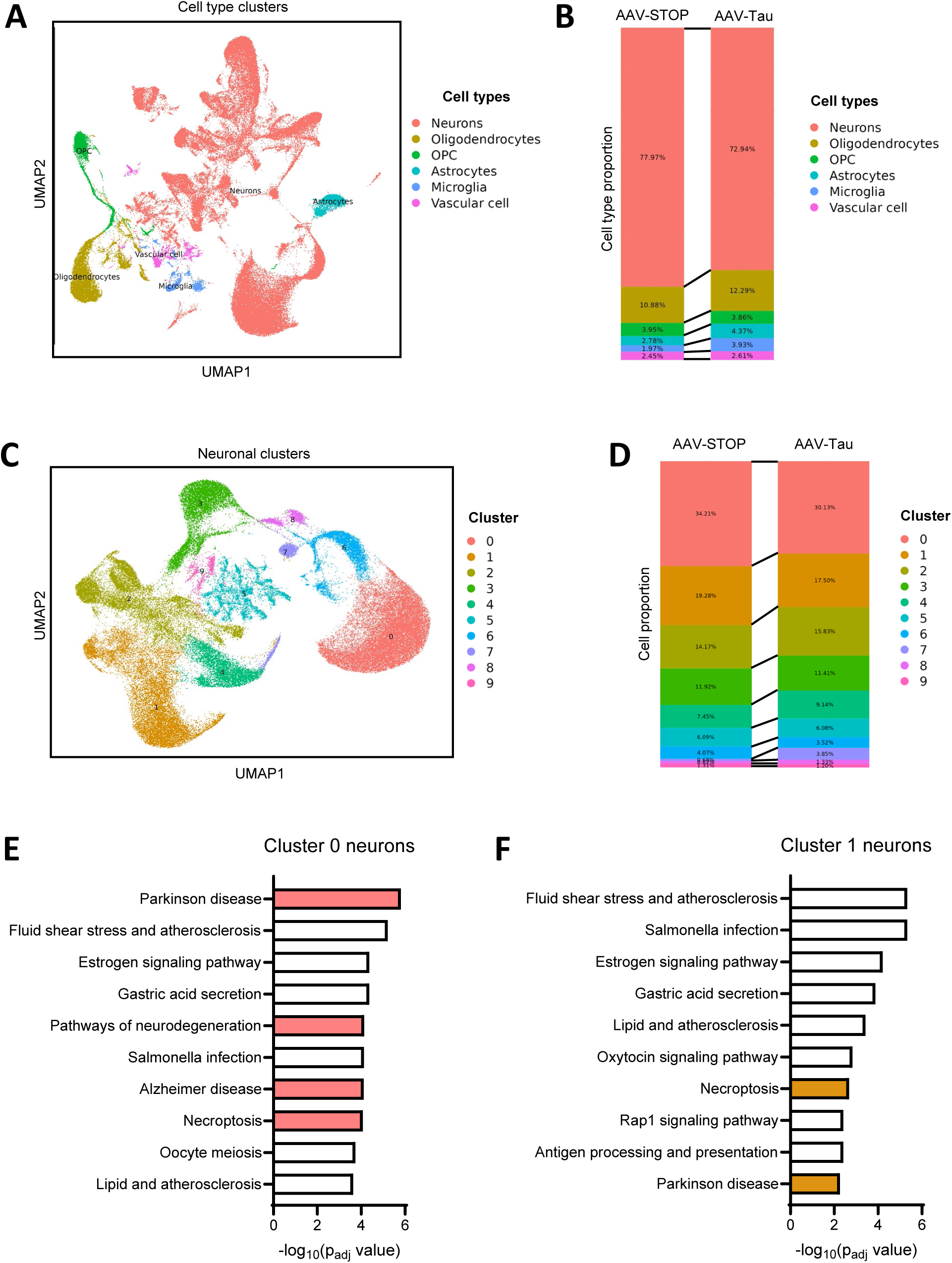
A) UMAP plot of single nuclei isolated from hippocampi of AAV-inoculated animals. Cells clustered into six groups based on genome wide expression patterns. B) Cell counts, expressed as a proportion, in hippocampi of 6-week-old animals injected with AAV-STOP and AAV-Tau. C) UMAP plot of single neuronal nuclei isolated from hippocampi of AAV-inoculated animals, showing 10 clusters of neurons. D) Cell counts of neuronal subtypes, expressed as a proportion, among the 10 neuron clusters identified in hippocampi of 6-week-old animals injected with AAV-STOP and AAV-Tau. E) Top 10 pathways (KEGG) most significantly regulated in cluster 0 neurons from AAV-Tau animals compared to AAV-STOP. F) Top 10 pathways (KEGG) most significantly regulated in cluster 1 neurons from AAV-Tau animals compared to AAV-STOP.

Remarkably, AAV-Tau hippocampus tissue displayed a significant reduction in the neuron cell population, by percent of total cells, when compared with AAV-STOP (Fig. 3B), a finding consistent with our structural and histological analyses (Fig. 2). We further interrogated the neuronal cells and came up with 10 subclusters (Fig. 3C). Based on marker gene profiles, they fell into four subtypes: excitatory neurons from CA1 (Ex_CA1), excitatory neurons from CA2/3, excitatory neurons from the dentate gyrus (Ex_DG), and inhibitory neurons (Fig. S2B). Populational analysis revealed selective loss of Cluster 0 and Cluster 1 neurons, corresponding to Ex_DG and Ex_CA1, respectively (Fig. 3D). Pathway analysis on the differentially expressed genes between these neurons showed remarkable upregulation of multiple disease-related pathways such as “Parkinson’s Disease,” “Alzheimer’s Disease”, and “Pathways of neurodegeneration,” as well as cell death process, such as “Necroptosis” in the affected neuronal subsets (Fig. 3E,F). In addition, several calmodulin genes, such as *Calm1*, *Calm2*, and *CamK1d*, were differentially affected by wild-type tau in these hippocampal neurons, implicating a profound effect of tau on neuronal calcium signaling (Fig. S2C,D). Thus, snRNA-seq analysis confirmed a severe loss of hippocampal neurons by wild-type tau and identified molecular processes relevant to neurodegeneration in the neuronal populations.

### Glial responses to non-mutated hTau

It is well known that tau pathology is associated with a neuroinflammatory response, manifesting in the brain as reactive gliosis (Johnson and Lukens, 2023; Parra Bravo et al., 2024). GFAP^+^ astrocytes showed subtle increases, most significantly in the CA1 field of the hippocampus, as early as 3 weeks post-inoculation with AAV-Tau (Fig. 4A). While the complement pathway is activated in various neurological diseases, the central component C3, a product of reactive astrocytes, has been functionally implicated in the pathogenesis of tauopathy models overexpressing mutant hTau (Litvinchuk et al., 2018; Wu et al., 2019). We found that hippocampal astrocytes displayed higher C3 expression in Tau-expressing brain at 6 weeks (Fig. 4B).

**Figure 4.**
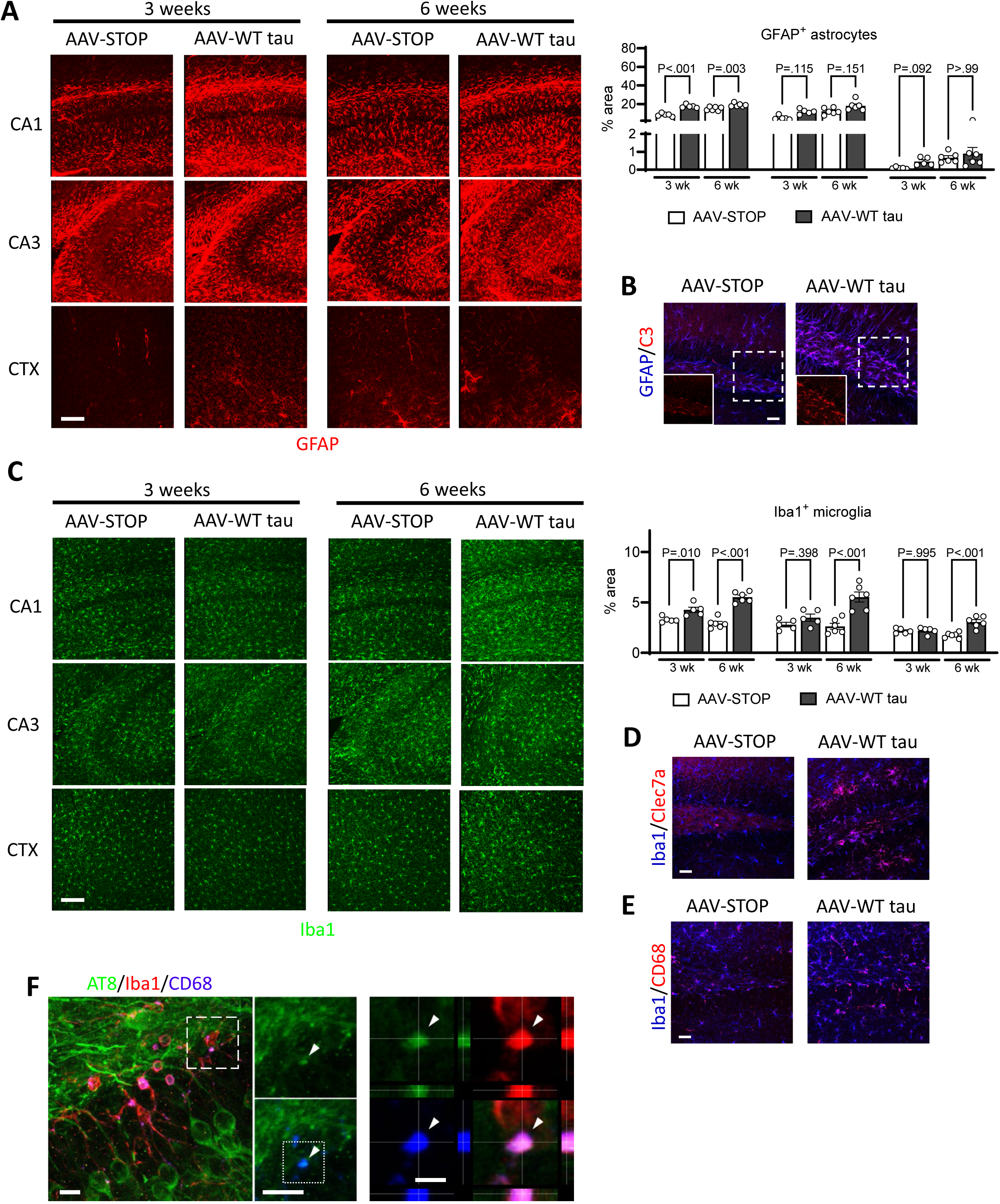
A) Representative confocal images and quantification of GFAP^+^ astrocytes in brain subregions at 3 and 6 weeks of age in AAV-Tau compared to AAV-STOP animals. Scale bar, 100 µm. B) Confocal images showing co-localized C3^+^ signal within GFAP^+^ astrocytes at 6 weeks of tau expression. Insets show isolated C3 channel. Scale bar, 10 µm. C) Representative confocal images and quantification of Iba1+ microglia in brain subregions at 3 and 6 weeks of age in AAV-Tau compared to AAV-STOP animals. Scale bar, 100 µm. D) Confocal images showing co-localized Clec7a^+^ signal within Iba1^+^ astrocytes at 6 weeks of tau expression. Scale bar, 10 µm. E) Confocal images showing co-localized CD68^+^ signal within Iba1^+^ astrocytes at 6 weeks of tau expression. Scale bar, 10 µm. F) A representative confocal image of several CA1 microglia expressing CD68^+^ lysosomal vesicles. AT8^+^ tau aggregates can be seen within these vesicles (arrowhead inside inset). The highlighted vesicle is further enlarged and presented as an orthogonal projection at the right. All scale bars, 10 µm.

Microglia are brain-resident immune phagocytes that fulfill a multitude of critical roles in the CNS (Paolicelli et al., 2022; Prinz et al., 2019). Fairly subtle increases in Iba1^+^ microglia were detected at 3 weeks of tau expression but more dramatic increases were evident by 6 weeks (Fig. 4C). Activated microglia are a feature of the brains of tauopathy models overexpressing mutant hTau (Johnson and Lukens, 2023; Parra Bravo et al., 2024). Likewise, we observed elevated expression of microglial activation markers, namely Clec7a (Fig. 4D) and CD68 (Fig. 4E), in the hippocampus by 6 weeks. We also found examples of AT8^+^ p-tau localized within CD68^+^ microglial vesicles (Fig. 4F), indicating a continuous engulfment of Tau by microglia. Our examination thus suggests an early astrocyte reaction, followed by microglial activation, in response to wild-type human tau expression in the rodent brain.

### Resistance to hTau-induced hippocampal neurodegeneration in mice lacking microglia

Depending on specific conditions, microglia activation may be reactive, protective, restorative, or pathogenic in the brain (Paolicelli et al., 2022). To examine the functional role of microglia in AD-relevant tauopathy, we adopted the *Csf1r^FIRE^* line (referred to here as “FIRE”), which completely lacks microglia throughout the lifespan due to the ablation of a super-enhancer element within the *Csf1r* gene (Fig. S3A) (Rojo et al., 2019). After inoculating AAV-Tau into FIRE mice, we observed a significant rescue of brain weight loss in FIRE mice (Fig. 5A), implying that microglia may be involved in neuronal deficits caused by Tau. Indeed, the neuronal density decreases seen in Tau-expressing wild-type animals, especially in the CA1, were abrogated in the brains of FIRE mice (Fig. 5B). These findings point to an essential role of microglia in mediating neurodegeneration induced by Tau.

**Figure 5.**
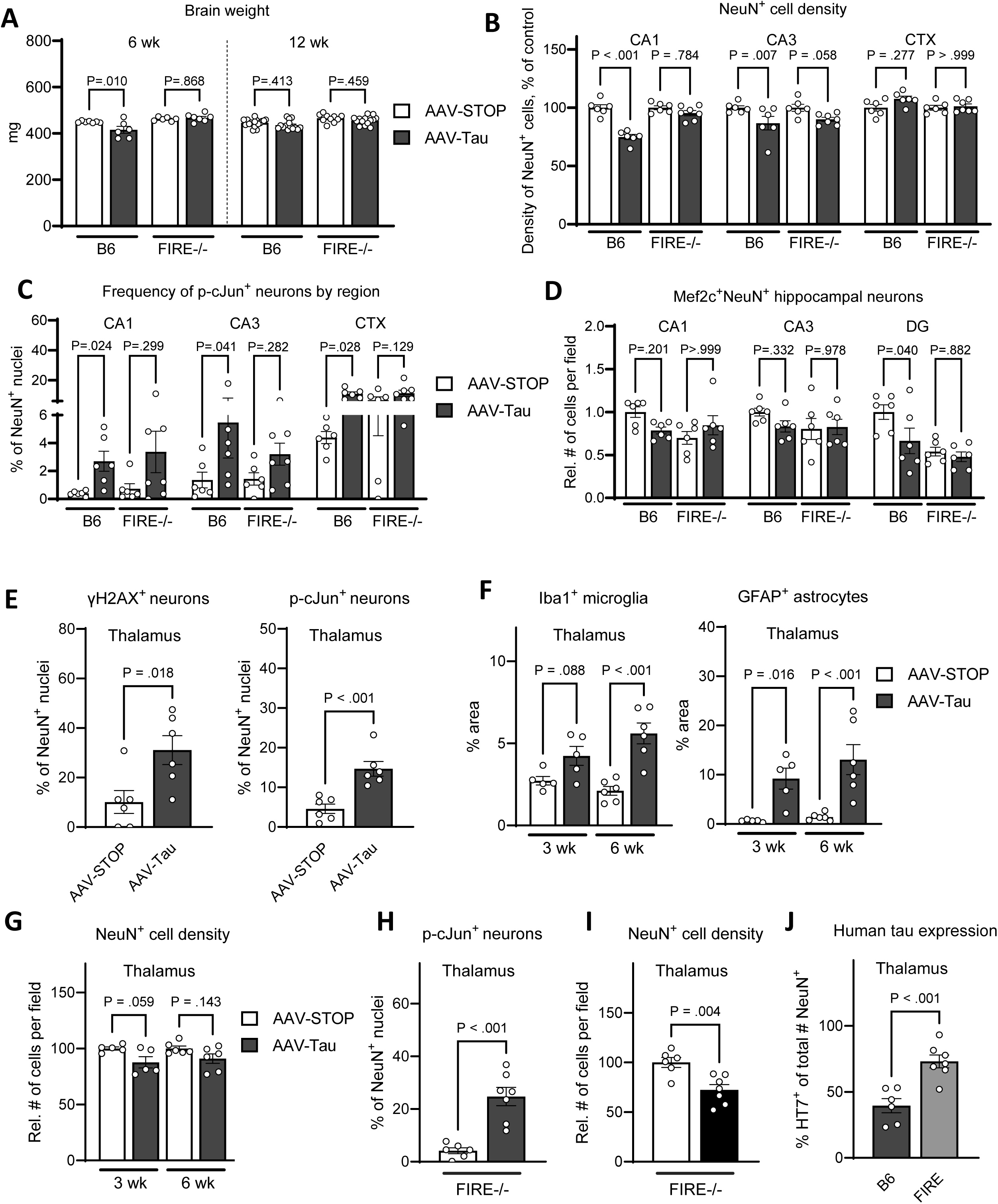
A) Quantification of brain weight at various time points in B6 and FIRE animals overexpressing wild-type tau compared to non-coding control vector. B) Quantification of relative neuronal density in B6 and FIRE animals overexpressing wild-type tau compared to non-coding control vector animals by region and age. C) Quantification of p-cJun^+^ neurons in brain subregions at 6 weeks of age in both B6 and FIRE mice expressing Tau or control vector. D) Quantification of the frequency of Mef2c^+^NeuN^+^ nuclei at 6 weeks of age in multiple regions from both B6 and FIRE mice expressing Tau or control vector. See Fig. S3B for representative images. E) Quantification of γH2AX^+^ and p-cJun^+^ neurons in the B6 thalamus at 6 weeks of age in AAV-Tau compared to AAV-STOP animals. F) Quantification of GFAP^+^ astrocytes and Iba1^+^ microglia in the B6 thalamus at 3 and 6 weeks of age in AAV-Tau compared to AAV-STOP animals. G) Quantification of relative thalamic neuronal density under AAV-Tau compared to AAV-STOP expression in B6 animals by age. See Fig. S3E for representative images. H,I) Quantification of relative thalamic neuronal density (H) and p-cJun^+^ neurons (I) at 6 weeks in FIRE animals expressing AAV-Tau compared to AAV-STOP. J) Quantification of HT7^+^ neuron percentage in thalamus of B6 and FIRE mice expressing AAV-Tau.

To assess the impact of microglial processes on tau-induced alterations, we examined a few other markers of neuronal processes, namely p-cJun, which is a key molecule in the DLK-MAPK stress pathway, and Mef2c, a master regulator of neuronal function that is associated with cognitive resilience in AD (Barker et al., 2021; Zhang and Zhao, 2022). While increased p-cJun was observed elevated in B6 animals expressing tau (Fig. 2E), despite having no microglia, FIRE mice expressing tau displayed similar levels of neuronal p-cJun (Fig. 5C). On the other hand, we observed a trend of mild reduction of hippocampal Mef2c^+^ neuron density in B6 animals expressing Tau (Fig. 5D, Fig. S3B). Maintained in a mixed C57BL/6:CBA background (Rojo et al., 2019), young FIRE mice held somewhat lower Mef2c^+^ neuronal densities in the hippocampus than their age-matched B6 counterparts (Fig. 5D). However, Tau expression did not affect Mef2c^+^ neuron density in FIRE mice (Fig. 5D). Taken altogether, these data suggest that microglia play an essential role in the hippocampal neurodegenerative process prompted by wild-type hTau.

### Tau- and microglia-modulated neurodegeneration in the thalamus

During examination, we noticed a particularly concentrated expression of transgenic Tau in the thalamus (Fig. 1B). In fact, whereas the percentage of neurons in other areas that expressed the transgene were normally under 20% (Fig. 1C), thalamic neurons expressed Tau at a level twice as frequently (Fig. S3C). Within the thalamus, Tau appeared to be expressed almost exclusively in excitatory neurons (marked by CAMKIIα), whereas inhibitory neurons (marked by PV) were devoid of human tau (Fig. S3D). Consistent with greater Tau expression, the frequency of neurons displaying DNA damage in the form of γH2AX expression, and those with elevated p-cJun, were both significantly elevated (Fig. 5E), surpassing other regions (Figs. 2D,E). In addition, GFAP^+^ astrocytes were far more drastically induced in the thalamus, while the level of Iba1^+^ microglia was largely comparable to other regions (Fig. 5F, Figs. 4A,C). However, no significant change of neuronal density was detected in the thalamus (Figs. 5G, S3E), a contrast to the hippocampal regions (Fig. 2C). Thus, thalamic neurons abundantly express Tau and display heightened stress response amidst significant gliosis, yet without undergoing neurodegeneration in wild-type mice.

Given the profound ongoing gliosis, we sought to determine the effects of microglia on Tau-induced cellular changes in the thalamus of FIRE mice. Surprisingly, thalamic neuronal cell density was significantly reduced in the absence of microglia (Fig. 5H). Accompanying this, the neuronal stress and injury marker p-cJun was elevated among the thalamic neurons in FIRE (∼25%, Fig. 5I) compared to wild-type animals (∼15%, Fig. 5E). Additionally, the overall population of Tau-expressing neurons was significantly increased in thalamus without microglia (Fig. 5J). Altogether, we have detected a prominent wild-type hTau expression in the thalamic neurons and uncovered an unexpected protective role of microglia in restraining Tau-mediated neurodegeneration in the thalamus.

## Discussion

By comparing neuronal expression of non-mutated versus FTLD-related human tau in the mouse brain, we have demonstrated a superior capacity of wild-type tau to induce neurodegeneration *in vivo*. The findings align with our previous observations from primary neurons and highlight an AD-relevant tau-mediated pathogenesis that has not been sufficiently recognized and investigated. Beyond neuronal intrinsic changes, our results further demonstrated that glial responses accompanied the course of tauopathy with wild-type hTau and revealed contrasting roles of microglia in mediating or restraining neurodegenerative processes from different brain regions.

Unlike primary tauopathies, AD patients possess higher levels of tau proteins in their brains without genetic mutations in the *MAPT* gene. However, rodents overexpressing FTLD-related tau mutations are commonly used to model clinical AD tau pathology (Qian et al., 2024). Our finding that wild-type hTau induces severe degenerative sequelae over mutant tau on the brain, especially the hippocampal region, is consistent with recent reports. Middle-aged mice expressing AAV-mediated wild-type hTau exhibited more profound brain atrophy than those expressing P301L mutant tau despite the latter accumulating higher tau protein aggregation (Tetlow et al., 2023). Further, a direct comparison of transgenic mice genetically matched in overexpressing wild-type versus P301L hTau has revealed a greater pathogenicity of wild-type hTau, which was accompanied by exaggerated tau hyperphosphorylation and early cognitive impairment (Gamache et al., 2020). Altogether, these studies highlight the importance to establish an AD-relevant tauopathy model and delineate the underlying mechanism that is likely disease-specific.

AD and other tauopathies affect distinct brain regions, which results in diverging clinical presentations. In AD, tau pathology is commonly believed to initiate in the entorhinal cortex, spread to limbic regions, which includes the hippocampus, and then migrate to the neocortex (Arnsten et al., 2021; Braak and Braak, 1991). Our wild-type hTau model displays not only neuronal tau expression in the hippocampus but also, together with signs of brain atrophy, selective neuronal loss in the hippocampal areas. Beyond the brain regions mentioned above, thalamic neuropathology has been implicated in early AD (Rüb et al., 2016; Yadav et al., 2024). Most recently, Sárkány et al. reported that tau filaments accumulated early and progressively inside the glutamatergic neurons at the anterodorsal thalamus of AD patients (Aggleton et al., 2016). Here, we have uncovered abundant hTau expression, enhanced stress response, and elevated gliosis in the thalamus, a regional pathology not present in transgenic models. As increasingly recognized, the thalamus critically participates in memory and cognitive functions via various network connections, such as the Papez circuit (Roy et al., 2022; Yadav et al., 2024). Therefore, our hTau model offers a unique opportunity to dissect various aspects of AD-relevant tau pathology.

In primary neuronal cultures, non-mutated hTau triggers neurotoxicity via synergistic MAPK-DLK signaling and DNA damage response (Li et al., 2024). Yet, characterization of *in vivo* neuronal wild-type hTau expression has revealed intriguing complex molecular and cellular events, both neuron-intrinsic and neuron-extrinsic. Wild-type hTau expression was sufficient to cause the accumulation of phosphorylated tau, together with MAPK-DLK signaling and DNA damage response in hippocampal neurons. Our snRNAseq analysis validated the selective hippocampal neuronal loss and revealed the enrichment of pathways related to neurodegenerative diseases in the mostly affected excitatory neuron populations. Tau upregulated the necroptosis pathway in both Ex_DG and Ex_CA1 neurons that were significantly diminished in numbers due to wild-type hTau expression. Of note, necroptosis was recently shown to play a central role in mediating the loss of human tangle^+^ neurons in an AD xenograft model (Balusu et al., 2023). Moreover, dysregulated neuronal calcium signaling by hTau is consistent with earlier findings that wild-type hTau induces memory deficits by calcineurin-mediated inactivation of nuclear CaMKIV/CREB signaling (Yin et al., 2016).

Glia cells are increasingly recognized to play crucial roles in a wide range of neurological diseases, especially neurodegeneration. In the hippocampus expressing wild-type hTau, we have detected an early astrocyte response followed by a subsequent microglial activation. Moreover, a robust glial reaction to wild-type hTau manifested in the thalamus. These findings echo the early and persistent glial responses in transgenic mouse models expressing mutant tau (Yoshiyama et al., 2007). By examining FIRE mice, we discovered that microglia seemingly exert contrasting regional functional influences: while wild-type hTau-mediated hippocampal neuronal loss decreased in the FIRE brain, neuronal loss only incurred in the thalamus in the absence of microglia. This process, in part, is comparable to mutant hTau-mediated hippocampal degeneration fostered by aberrantly activated microglia (Johnson and Lukens, 2023; Shi et al., 2017; Udeochu et al., 2023).

However, microglia’s role in brain pathophysiology is highly complex. In the lifelong absence of microglia, FIRE mice offer a novel and valuable tool to investigate the function of these brain immune cells. Intriguingly, spontaneous age-related thalamic degeneration, such as atrophy and neuronal loss, has been a prominent feature of aged FIRE mice (Munro et al., 2024). Crossed with a xenotolerant strain, adult FIRE mice also display a selective pathology in the thalamus (Chadarevian et al., 2024). Comparatively, the FIRE mice we examined were fairly young – up to 6 weeks old. Given the high Tau load and stress response in our model, these observations indicate a key protective function of thalamic microglia in maintaining the viability of resident neurons, which otherwise would be particularly vulnerable to neuropathological changes during brain aging or AD.

Taken together, we have established an AD-relevant tauopathy model in young mice and uncovered neuron-intrinsic and -extrinsic responses associated with wild-type hTau-mediated neurodegeneration. These findings may inspire further research on the regional functions of microglia and the delineation of protective versus pathogenic roles of brain glia cells.

## Acknowledgments

This research was funded by the National Institute on Aging, National Institutes of Health, grant numbers AG057587 and AG074283; National Heart, Lung, and Blood Institute, National Institutes of Health, grant number HL154720-03S1; National Institute of Diabetes and Digestive and Kidney Diseases, National Institutes of Health, grant number DK122708-03S1; BrightFocus ADR A20183775; Brown Foundation 2020 Healthy Aging Initiative; and Pilot Grant support from UTHealth McGovern Medical School (W.C.). The content is solely the responsibility of the authors and does not necessarily represent the official views of the National Institutes of Health. We extend our gratitude to Dr. Mathew Blurton-Jones from the University of California, Irvine, for generously providing live FIRE mice, Mingyi Lv for assistance with RNAseq analysis, and Dr. Michael Beierlein at UTHealth for advising thalamus biology. We thank the technical support from the Cancer Prevention and Research Institute of Texas (CPRIT RP240610), esp. the work by Dr. Xian Chen. The schematic was created with BioRender.

## Author contributions

Conceptualization, E.R.R and W.C.; Methodology, E.R.R, Q.W, S.L, and S.H.; Formal analysis, E.R.R, Q.W, K.H, M.L, and E.E.; Resources: J.J.H, W.L, C.P, and X.Z.; Writing – original draft, E.R.R and W.C.; Writing – review & editing, E.R.R. and W.C.; Visualization, E.R.R, Q.W, and K.H.; Validation, Y.F. and C.J.; Supervision, W.C.; Project administration, W.C.; Funding acquisition, W.C. All authors have read and agreed to the manuscript.

## EXPERIMENTAL MODEL AND SUBJECT DETAILS

### Mice

C57BL/6J mice were obtained from the National Institute on Aging (NIA). FIRE mice in a mixed C57B/6:CBA background were generated by Clare Pridans (Rojo et al., 2019). Mice with *ad libitum* access to food and water and were housed in mixed-genotype groups of 3-5 per cage under specific pathogen-free conditions and standard light/dark cycle. Both male and female mice were used in experiments. Mice were analyzed at specific timepoints as noted in the study, and precise ages of all animals used in experiments are listed in respective figure legends. All animal procedures were performed in accordance with NIH guidelines and with the approval of the Institutional Animal Care and Use Committee at University of Texas Health Science Center-Houston.

### AAV delivery

The generation and preparation of all AAV vectors used in this study were described previously(Li et al., 2024). For *in vivo* inoculation, frozen stock solutions of AAV particles were thawed and diluted with vehicle composed of sterile PBS with 10% trypan blue (for visual confirmation of correct injection) to the appropriate concentration and kept on ice. Neonatal mouse pups were chilled on an ice-cold aluminum block until fully anesthetized. AAVs were delivered into the intracerebroventricular spaces of neonatal mouse pups (on p0, the day of birth) using a Hamilton Gas-tight syringe (Hamilton Company, Reno, NV) fitted with a steel 32-gauge needle (point style 4 with a 12° bevel). We used a dose of 5 x 10^9^ gc in 2 µL vehicle into each hemisphere of the brain, for a total dose of 1 x 10^10^ gc in 4 µL vehicle per animal. Injected neonates were allowed to regain body temperature on a heating pad before being placed back into parent cages.

### Volumetric MRI procedure

*In vivo* MRI was performed on a 7T Bruker BioSpec MRI system (Bruker BioSpin, Billerica, MA) at the MRI core facility of UTHealth Houston. Specifically, T_2_-weighted images were acquired using the RARE sequence with the following imaging parameters: TR/TE: 6000/60 ms, in-plane resolution: 100 x 100 μm^2^, slice thickness: 0.5 mm, and a total of 30 slices to cover the whole brain. Brain structures of interest were manually segmented using ITK-SNAP software (version 3.8.0) (Yushkevich et al., 2006). Briefly, appropriate structures were delineated on each 2D image slice in which the structure appeared, and 3D volumes were calculated based on this segmentation, which yielded a total 3D volume in mm^3^ for each structure.

## METHOD DETAILS

### Single-nuclei RNA sequencing

The mouse hippocampus was dissected and snap-frozen at -80°C. Tissue (100 mg) was powdered on dry ice using a scalpel and transferred to a 1.5 mL microtube. Homogenization was performed by adding 0.5 mL of freshly prepared Homogenization Buffer (320 mM Sucrose, 5 mM CaCl_2_, 3 mM Mg(Ac)_2_, 10 mM Tris pH 7.8, 0.1 mM EDTA, 0.1% NP40, 0.1 mM Protease inhibitor, 1 mM Beta-mercaptoethanol, 1 U/μL RNase inhibitor), followed by gentle pipetting and incubation on ice for 10 min. The homogenate was passed through a 30 μm strainer and centrifuged at 300 g for 3 min at 4°C. The pellet was resuspended in 0.5 mL Homogenization Buffer and mixed with 0.5 mL Upper Gradient Centrifugation Buffer (50% OptiPrep Gradient, 5 mM CaCl_2_, 3 mM Mg(Ac)_2_, 10 mM Tris pH 7.8, 0.1 mM Protease inhibitor, 1 mM Beta-mercaptoethanol, 1 U/μL RNase inhibitor). The mixture was gently layered on top of 1 mL of Lower Gradient Centrifugation Buffer (29% OptiPrep Gradient, 1 U/μL RNase inhibitor) in a 2 mL microtube.

The gradient was centrifuged at 7,200 g for 15 min at 4°C. The supernatant was carefully discarded. The nuclei pellet was resuspended in the PBS containing 0.04% BSA and 1 U/μL RNase inhibitor to achieve a concentration of 700-1200 nuclei/μL.

Nuclei (∼20,000 per sample) were loaded onto a 10x Genomics Next GEM chip G. Libraries were prepared using the Chromium Next GEM Single Cell 3ʹ Library and Gel Bead kit v3.1 (10x Genomics), following the manufacturer’s protocol, and sequenced on an Illumina NovaSeq 6000. The sequencing data was generated by the UTHealth Houston Cancer Genomics Core.

### Single-cell RNA preprocessing and cell type annotation

The raw scRNA-seq data were processed using CellRanger v4.0, which performed initial data alignment, filtering, and UMI counting. The mouse reference genome (mm10) was utilized for accurate mapping of sequencing reads to genes. After preprocessing with CellRanger, the resulting gene-barcode matrices were imported into R for downstream analyses. The quality control and preprocessing steps were performed using Seurat (v4.4.0), and batch effect removal was performed using Harmony(Butler et al., 2018; Korsunsky et al., 2019). To preprocess the scRNA-seq data, we applied stringent quality control thresholds to ensure the reliability and biological relevance of the downstream analysis. Cells were filtered based on the following criteria: a minimum of 200 detected genes (nFeature_RNA > 200) to exclude low-quality or empty droplets, and a total RNA count between 1,000 and 20,000 (1000 < nCount_RNA < 20000) to eliminate cells with extremely low or high RNA content, which could indicate technical artifacts or doublets. Additionally, cells with a mitochondrial gene percentage exceeding 5% (percent.mt < 5) were excluded to remove cells with potential stress or apoptosis signals.

Cell clustering for the scRNA-seq data was performed using the Seurat package in R, utilizing normalized gene expression profiles to identify distinct cell populations. The resolution was set as 0.1 for the clustering. Cell type annotation was carried out using the UCell package, which computes gene set enrichment scores based on marker genes curated from established literature and databases (Andreatta and Carmona, 2021). Each cluster was assigned to a corresponding cell type based on the highest enrichment score.

### Differential gene expression analysis and enrichment analysis for each cell type

We performed differential expression analysis to identify aging-related genes across different mouse types (i.e., AAV-Tau vs. AAV-STOP) using the ‘FindMarkers’ function in the Seurat package for each cell type. Differentially expressed genes (DEGs) were defined as those with an adjusted p-value < 0.05 and |logFC| greater than the threshold calculated as mean(abs(logFC)) + 2 × sd(abs(logFC)).

To investigate the potential biological roles of these DEGs, we performed KEGG pathway enrichment analysis using the clusterProfiler v4.10.1 package(Yu et al., 2012). Separate analyses were conducted for upregulated and downregulated DEGs within each cell type. Only pathways with an adjusted p-value < 0.05 were considered significant.

### Identification of subpopulation for major cell types

After the cell type annotation for the major cell types, we extract the certain cell type for the subpopulation analysis. Similarly with the major cell type annotation, Seurat and UCell packages were used in this step(Andreatta and Carmona, 2021; Butler et al., 2018).

### Single-cell trajectory reconstruction and analysis

Single-cell pseudotime trajectories were constructed with Monocle (version 2.30.1) (Qiu et al., 2017). Briefly, we first selected the cell type and the subpopulations with their gene expression matrix. Monocle applies reversed graph embedding, a machine learning approach, to construct a parsimonious principal graph that reduces high-dimensional expression profiles into a low-dimensional space. Each cell is projected onto this space and organized into a trajectory with distinct branch points with a pseudotime value.

### Immunofluorescence

Mice were perfused with ice-cold saline (0.9% NaCl) after deep anesthesia with ketamine/xylazine, and brains were extracted, fixed overnight at 4°C in 4% paraformaldehyde (Santa Cruz, cat# sc-281692), and dehydrated in 30% sucrose until sectioning. Brains were sectioned into 30 μm tissue sections using a freezing microtome, and tissue was stored in cryoprotectant at -20°C. For staining, floating sections were washed in phosphate-buffered saline (PBS) and blocked for 1 h at RT in a blocking buffer of 10% normal donkey serum (Millipore, cat# S30-100ML) and 1% Triton X-100 in TBS. Primary antibodies were diluted in blocking buffer and applied to sections overnight at 4°C. Tissues were washed with PBS-T (PBS with 0.1% Tween-20) three times, then incubated with fluorescent secondary antibodies diluted in blocking buffer for 1 h at RT. After final washing in PBS-T, sections were mounted on glass slides, allowed to dry, and coverslipped with ProLong Glass Antifade mountant (Life Technologies, cat# P36982).

### Image quantification

All imaging in this study was performed with a Leica SP8 system. For all quantitative analyses in this study, multiple images were used as technical replicates from each animal. For counting of neurons, NeuN antibody was applied to tissues to visualize individual neuronal nuclei. In some experiments, other markers were co-labelled with NeuN, such as p-cJun or Mef2c. Single-plane confocal images from regions of interest were subjected to manual counting in the Leica confocal software, and data were recorded for total NeuN^+^ cells per field, and those with and without co-expression of markers pertinent to the experiment.

For area fraction calculation, Z-stacks were converted to maximum intensity projections and loaded into ImageJ (NIH) for analysis. Images were thresholded manually to overlap fluorescence signals, and percent area was measured. For assessment of hippocampal layer thickness, confocal images of NeuN^+^ cell layers (CA1 and CA3) were acquired using a 20X lens. For each field of view, the thickness of the layer at three to five points was collected and averaged, and multiple sections were imaged per animal.

### Transmission electron microscopy (TEM)

Hippocampal tissue was imaged following standard electron microscopy procedures using a Ted Pella Bio Wave processing microwave with vacuum attachments. Mice were perfused using a saline flush followed by Modified Karnovski’s fixative in sodium cacodylate buffer for 40 minutes per animal. Whole heads were dissected and brains were sectioned at 1mm thickness, followed by micro-dissection of hippocampal tissues. The tissue was covered in 2% paraformaldehyde, 2.5% glutaraldehyde, in 0.1 M sodium cacodylate buffer at pH 7.4. After dissection the tissues were incubated in the fixative in a cold room rotator for three days. The pre-fixed tissue sections were then fixed again, followed by 3x Millipore water rinses, post-fixed with 1% aqueous osmium tetroxide, and rinsed again 3x with Millipore water. Concentrations from 25%-100% of ethanol were used for the initial dehydration series, followed with propylene oxide as the final dehydrant. Samples were gradually infiltrated with 3 ratios of propylene oxide and Embed 812, finally going into 3 changes of pure resin under vacuum. Samples were allowed to infiltrate in pure resin overnight on a rotator. The samples were embedded into appropriate molds and cured in the oven at 62°C for five days. The polymerized samples were thinly sectioned at 48-50 nm in a Leica UC7 Ultramicrotome. The grids were stained with 1% uranyl acetate for ten minutes followed by lead citrate for two minutes before TEM examination. Grids were viewed in a JEOL JEM 1400 Plus transmission electron microscope at 80kV. Images were captured using an AMT XR-16 mid-mount 16 mega-pixel digital camera.

## QUANTIFICATION AND STATISTICAL ANALYSIS

All data in bar plots are presented as means ± s.e.m. GraphPad Prism (v10.0.0) was used for all statistical analyses and graph plotting. All data was tested for normality using the Shapiro-Wilk test. For normally distributed data, differences between two groups were analyzed by two-tailed Student’s t-tests (with Holm-Šídák’s post-hoc multiple comparisons tests, where applicable), and differences between three or more groups were analyzed by one-way ANOVA with Šídák’s post-hoc multiple-comparisons tests. For comparisons including non-normally distributed data, differences between two groups were analyzed by Mann-Whitney U Tests (with Holm-Šídák’s post-hoc multiple comparisons tests, where applicable), and differences between three or more groups were analyzed by Kruskal-Wallis tests with Dunn’s post-hoc multiple-comparisons tests. P or P_adj_ values less than 0.05 were considered significant, and those over 0.05 were considered non-significant. All exact P values are listed within plots. Female and male mice were used for all experiments. All micrographs shown are images representative of multiple replicates as noted in legends. No statistical methods were used to predetermine sample sizes, but similar publications were used as guidelines. Biorender.com was used to generate schematics within the figures.

---

**Figure S1.**
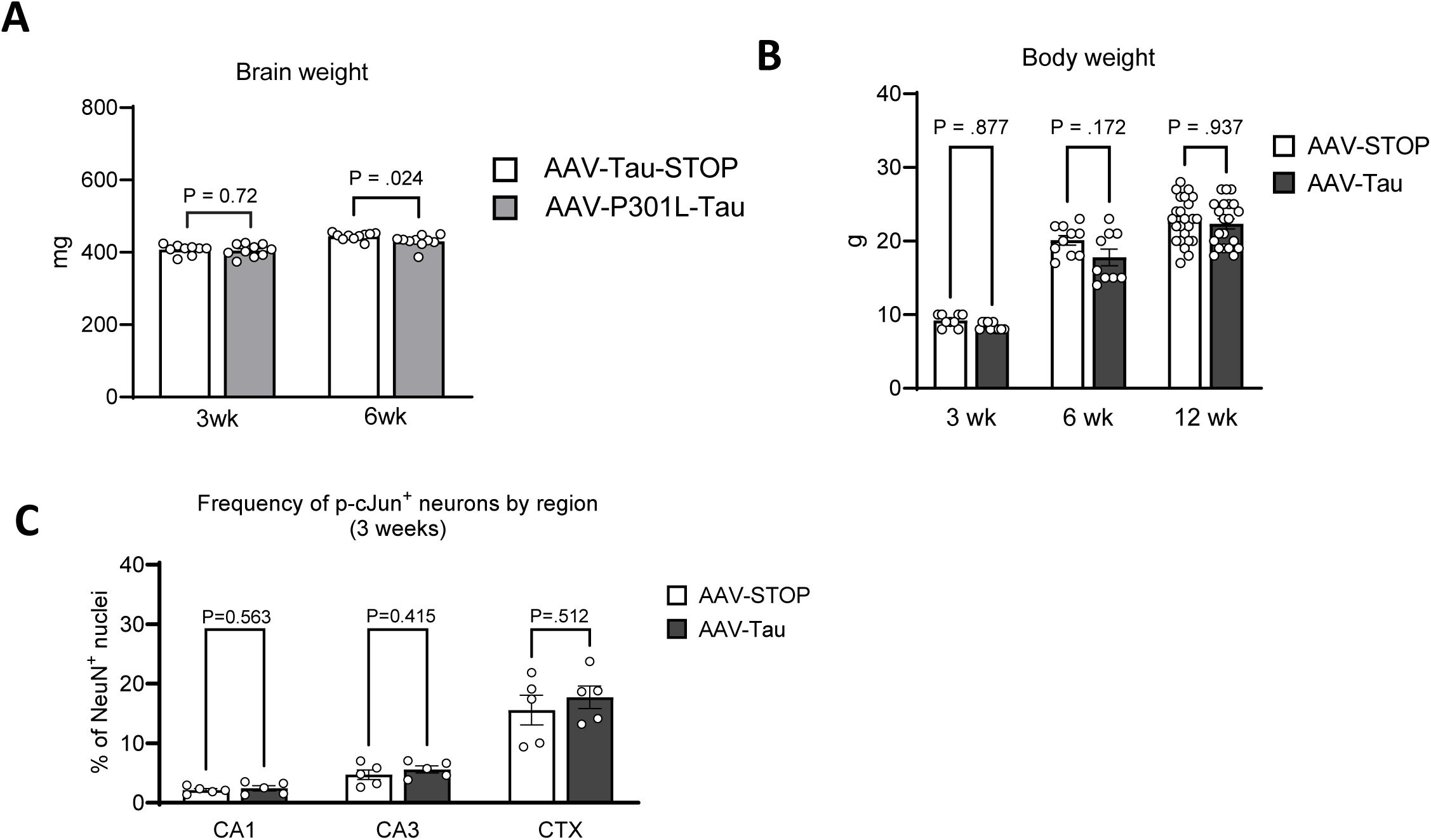
A) Quantification of brain weight at 3- and 6-weeks post-inoculation in animals overexpressing P301L mutant tau compared to non-coding control vector. B) Quantification of body weight at 3-, 6-, and 12-weeks post-inoculation in animals overexpressing wild-type hTau compared to non-coding control vector. C) Quantification of p-cJun^+^ neurons in brain subregions at 3 weeks of age in AAV-Tau compared to AAV-STOP animals.

**Figure S2.**
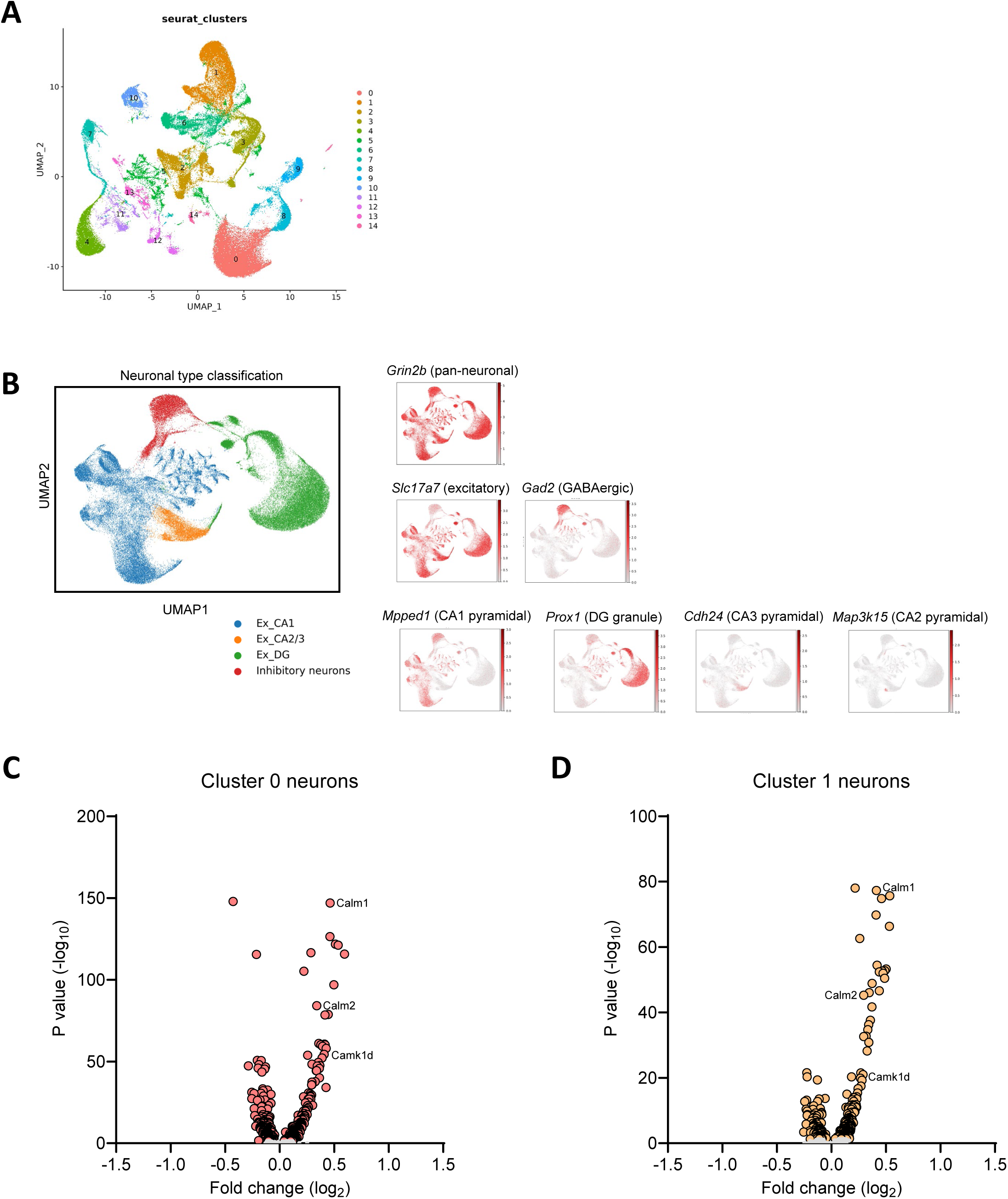
A) UMAP plots showing annotation of neuronal clusters with neuron type markers. B) Classification of major neuronal types, including excitatory cells from CA1, CA2/3 and DG, and inhibitory cells. C,D) Volcano plot of gene expression changes in cluster 0 (C) and cluster 1 (D) neurons in 6-week-old AAV-Tau animals compared to AAV-STOP animals, with significant DEGs colored.

**Figure S3.**
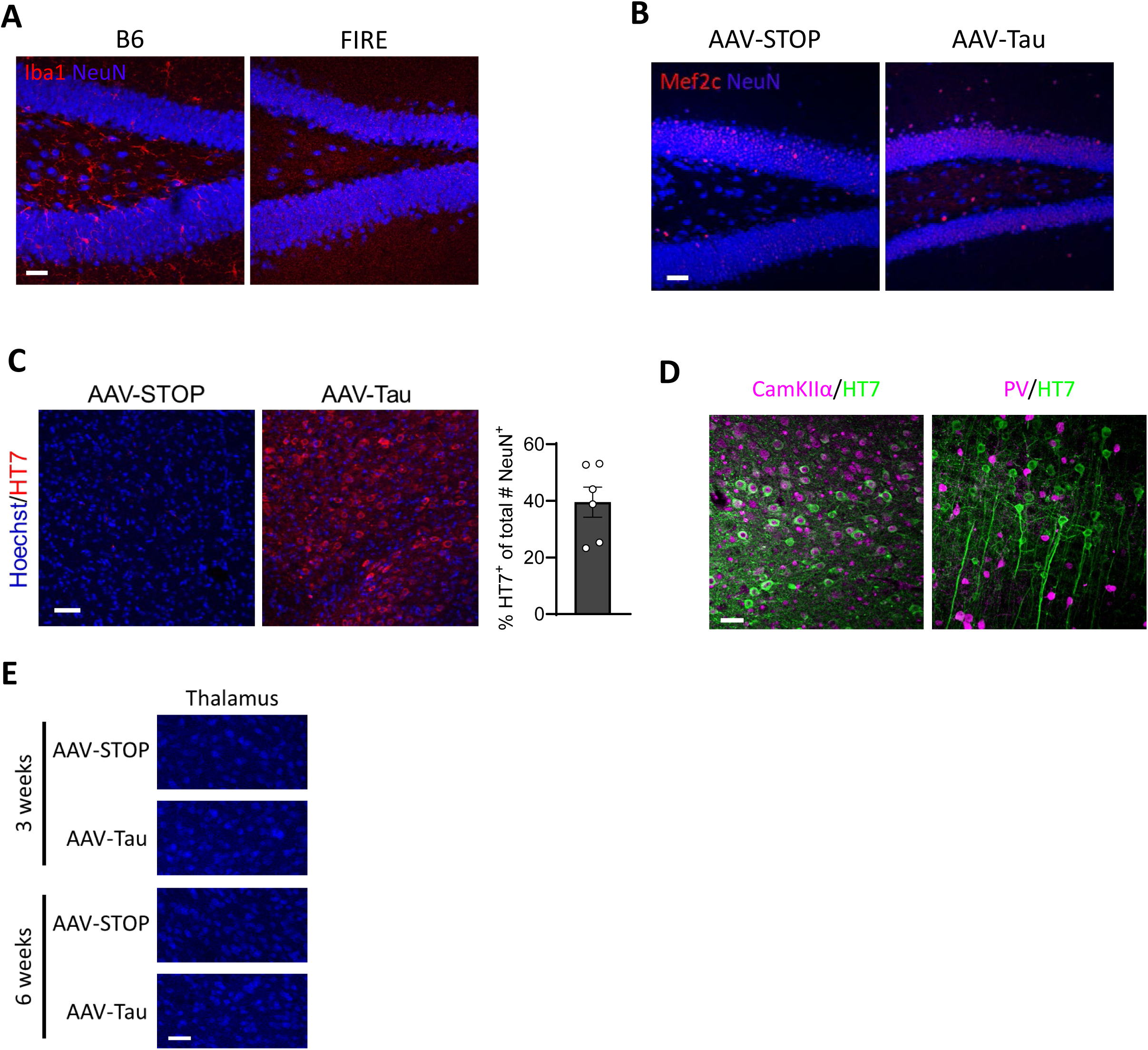
A) Representative confocal images of Iba1 staining in the dentate gyrus of B6 wild-type mice and FIRE mice. Scale bar, 50 µm. B) Representative confocal images of dentate gyrus expression of Mef2c in neuronal nuclei. Scale bar, 50 µm. C) Representative confocal images of thalamus in AAV-Tau and AAV-STOP animals at 6 weeks of age. Scale bar, 50 µm. Quantification of % HT7^+^ neurons in the thalamus of 6-week-old AAV-Tau animals. D) Distribution of transgenic human tau (HT7) among neuronal subtypes, including excitatory (CamKIIα^+^) and inhibitory (PV^+^) neurons in the cortex and thalamus. Scale bar, 30 µm. E) Representative confocal images of NeuN^+^ neurons in thalamus, at two time points after neonatal inoculation with AAV-STOP or AAV-Tau. Scale bar, 50 µm.

## References

Aggleton, J.P., A. Pralus, A.J. Nelson, and M. Hornberger. 2016. Thalamic pathology and memory loss in early Alzheimer’s disease: moving the focus from the medial temporal lobe to Papez circuit. Brain 139:1877–1890.

Andreatta, M., and S.J. Carmona. 2021. UCell: Robust and scalable single-cell gene signature scoring. Computational and structural biotechnology journal 19:3796–3798.

Apostolova, L.G., A.E. Green, S. Babakchanian, K.S. Hwang, Y.Y. Chou, A.W. Toga, and P.M. Thompson. 2012. Hippocampal atrophy and ventricular enlargement in normal aging, mild cognitive impairment (MCI), and Alzheimer Disease. Alzheimer Dis Assoc Disord 26:17–27.

Arnsten, A.F.T., D. Datta, K. Del Tredici, and H. Braak. 2021. Hypothesis: Tau pathology is an initiating factor in sporadic Alzheimer’s disease. Alzheimer’s & dementia: the journal of the Alzheimer’s Association 17:115–124.

Balusu, S., K. Horré, N. Thrupp, K. Craessaerts, A. Snellinx, L. Serneels, D. T’Syen, I. Chrysidou, A.M. Arranz, A. Sierksma, J. Simrén, T.K. Karikari, H. Zetterberg, W.-T. Chen, D.R. Thal, E. Salta, M. Fiers, and B. De Strooper. 2023. MEG3 activates necroptosis in human neuron xenografts modeling Alzheimer’s disease. Science 381:1176–1182.

Barker, S.J., R.M. Raju, N.E.P. Milman, J. Wang, J. Davila-Velderrain, F. Gunter-Rahman, C.C. Parro, P.L. Bozzelli, F. Abdurrob, K. Abdelaal, D.A. Bennett, M. Kellis, and L.H. Tsai. 2021. MEF2 is a key regulator of cognitive potential and confers resilience to neurodegeneration. Sci Transl Med 13:eabd7695.

Bohmann, D., T.J. Bos, A. Admon, T. Nishimura, P.K. Vogt, and R. Tjian. 1987. Human proto-oncogene c-jun encodes a DNA binding protein with structural and functional properties of transcription factor AP-1. Science 238:1386–1392.

Braak, H., and E. Braak. 1991. Neuropathological stageing of Alzheimer-related changes. Acta Neuropathol 82:239–259.

Butler, A., P. Hoffman, P. Smibert, E. Papalexi, and R. Satija. 2018. Integrating single-cell transcriptomic data across different conditions, technologies, and species. Nature biotechnology 36:411–420.

Chadarevian, J.P., J. Hasselmann, A. Lahian, J.K. Capocchi, A. Escobar, T.E. Lim, L. Le, C. Tu, J. Nguyen, S. Kiani Shabestari, W. Carlen-Jones, S. Gandhi, G. Bu, D.A. Hume, C. Pridans, Z.K. Wszolek, R.C. Spitale, H. Davtyan, and M. Blurton-Jones. 2024. Therapeutic potential of human microglia transplantation in a chimeric model of CSF1R-related leukoencephalopathy. Neuron 112:2686–2707 e2688.

Chang, C.W., E. Shao, and L. Mucke. 2021. Tau: Enabler of diverse brain disorders and target of rapidly evolving therapeutic strategies. Science 371:eabb8255.

Dujardin, S., C. Commins, A. Lathuiliere, P. Beerepoot, A.R. Fernandes, T.V. Kamath, M.B. De Los Santos, N. Klickstein, D.L. Corjuc, B.T. Corjuc, P.M. Dooley, A. Viode, D.H. Oakley, B.D. Moore, K. Mullin, D. Jean-Gilles, R. Clark, K. Atchison, R. Moore, L.B. Chibnik, R.E. Tanzi, M.P. Frosch, A. Serrano-Pozo, F. Elwood, J.A. Steen, M.E. Kennedy, and B.T. Hyman. 2020. Tau molecular diversity contributes to clinical heterogeneity in Alzheimer’s disease. Nat. Med. 26:1256–1263.

Gamache, J.E., L. Kemper, E. Steuer, K. Leinonen-Wright, J.M. Choquette, C. Hlynialuk, K. Benzow, K.A. Vossel, W. Xia, M.D. Koob, and K.H. Ashe. 2020. Developmental Pathogenicity of 4-Repeat Human Tau Is Lost with the P301L Mutation in Genetically Matched Tau-Transgenic Mice. J. Neurosci. 40:220–236.

Goedert, M., and R. Jakes. 2005. Mutations causing neurodegenerative tauopathies. Biochim. Biophys. Acta 1739:240–250.

Guo, T., W. Noble, and D.P. Hanger. 2017. Roles of tau protein in health and disease. Acta Neuropathol 133:665–704.

Hu, Y.Y., S.S. He, X. Wang, Q.H. Duan, I. Grundke-Iqbal, K. Iqbal, and J. Wang. 2002. Levels of nonphosphorylated and phosphorylated tau in cerebrospinal fluid of Alzheimer’s disease patients: an ultrasensitive bienzyme-substrate-recycle enzyme-linked immunosorbent assay. Am J Pathol 160:1269–1278.

Johnson, A.M., and J.R. Lukens. 2023. The innate immune response in tauopathies. Eur. J. Immunol. 53:e2250266.

Khatoon, S., I. Grundke-Iqbal, and K. Iqbal. 1992. Brain levels of microtubule-associated protein tau are elevated in Alzheimer’s disease: a radioimmuno-slot-blot assay for nanograms of the protein. J. Neurochem. 59:750–753.

Knopman, D.S., H. Amieva, R.C. Petersen, G. Chetelat, D.M. Holtzman, B.T. Hyman, R.A. Nixon, and D.T. Jones. 2021. Alzheimer disease. Nat Rev Dis Primers 7:33.

Korsunsky, I., N. Millard, J. Fan, K. Slowikowski, F. Zhang, K. Wei, Y. Baglaenko, M. Brenner, P.-r. Loh, and S. Raychaudhuri. 2019. Fast, sensitive and accurate integration of single-cell data with Harmony. Nature methods 16:1289–1296.

Koutsodendris, N., J. Blumenfeld, A. Agrawal, M. Traglia, O. Yip, A. Rao, M.J. Kim, M.R. Nelson, Y.-H. Wang, and B. Grone. 2023. APOE4-promoted gliosis and degeneration in tauopathy are ameliorated by pharmacological inhibition of HMGB1 release. Cell reports 42:

Li, S., E.R. Roy, Y. Wang, T. Watkins, and W. Cao. 2024. DLK-MAPK Signaling Coupled with DNA Damage Promotes Intrinsic Neurotoxicity Associated with Non-Mutated Tau. Mol. Neurobiol. 61:2978–2995.

Litvinchuk, A., Y.W. Wan, D.B. Swartzlander, F. Chen, A. Cole, N.E. Propson, Q. Wang, B. Zhang, Z. Liu, and H. Zheng. 2018. Complement C3aR Inactivation Attenuates Tau Pathology and Reverses an Immune Network Deregulated in Tauopathy Models and Alzheimer’s Disease. Neuron 100:1337–1353 e1335.

Mah, L.J., A. El-Osta, and T.C. Karagiannis. 2010. gammaH2AX: a sensitive molecular marker of DNA damage and repair. Leukemia 24:679–686.

Munro, D.A.D., N. Bestard-Cuche, C. McQuaid, A. Chagnot, S.K. Shabestari, J.P. Chadarevian, U. Maheshwari, S. Szymkowiak, K. Morris, M. Mohammad, A. Corsinotti, B. Bradford, N. Mabbott, R.J. Lennen, M.A. Jansen, C. Pridans, B.W. McColl, A. Keller, M. Blurton-Jones, A. Montagne, A. Williams, and J. Priller. 2024. Microglia protect against age-associated brain pathologies. Neuron 112:2732–2748 e2738.

Paolicelli, R.C., A. Sierra, B. Stevens, M.E. Tremblay, A. Aguzzi, B. Ajami, I. Amit, E. Audinat, I. Bechmann, M. Bennett, F. Bennett, A. Bessis, K. Biber, S. Bilbo, M. Blurton-Jones, E. Boddeke, D. Brites, B. Brone, G.C. Brown, O. Butovsky, M.J. Carson, B. Castellano, M. Colonna, S.A. Cowley, C. Cunningham, D. Davalos, P.L. De Jager, B. de Strooper, A. Denes, B.J.L. Eggen, U. Eyo, E. Galea, S. Garel, F. Ginhoux, C.K. Glass, O. Gokce, D. Gomez-Nicola, B. Gonzalez, S. Gordon, M.B. Graeber, A.D. Greenhalgh, P. Gressens, M. Greter, D.H. Gutmann, C. Haass, M.T. Heneka, F.L. Heppner, S. Hong, D.A. Hume, S. Jung, H. Kettenmann, J. Kipnis, R. Koyama, G. Lemke, M. Lynch, A. Majewska, M. Malcangio, T. Malm, R. Mancuso, T. Masuda, M. Matteoli, B.W. McColl, V.E. Miron, A.V. Molofsky, M. Monje, E. Mracsko, A. Nadjar, J.J. Neher, U. Neniskyte, H. Neumann, M. Noda, B. Peng, F. Peri, V.H. Perry, P.G. Popovich, C. Pridans, J. Priller, M. Prinz, D. Ragozzino, R.M. Ransohoff, M.W. Salter, A. Schaefer, D.P. Schafer, M. Schwartz, M. Simons, C.J. Smith, W.J. Streit, T.L. Tay, L.H. Tsai, A. Verkhratsky, R. von Bernhardi, H. Wake, V. Wittamer, S.A. Wolf, L.J. Wu, and T. Wyss-Coray. 2022. Microglia states and nomenclature: A field at its crossroads. Neuron 110:3458–3483.

Parra Bravo, C., S.A. Naguib, and L. Gan. 2024. Cellular and pathological functions of tau. Nat. Rev. Mol. Cell Biol.

Prinz, M., S. Jung, and J. Priller. 2019. Microglia Biology: One Century of Evolving Concepts. Cell 179:292–311.

Qian, Z., Y. Li, and K. Ye. 2024. Advancements and challenges in mouse models of Alzheimer’s disease. Trends Mol. Med.

Qiu, X., A. Hill, J. Packer, D. Lin, Y.-A. Ma, and C. Trapnell. 2017. Single-cell mRNA quantification and differential analysis with Census. Nature methods 14:309–315.

Rojo, R., A. Raper, D.D. Ozdemir, L. Lefevre, K. Grabert, E. Wollscheid-Lengeling, B. Bradford, M. Caruso, I. Gazova, A. Sanchez, Z.M. Lisowski, J. Alves, I. Molina-Gonzalez, H. Davtyan, R.J. Lodge, J.D. Glover, R. Wallace, D.A.D. Munro, E. David, I. Amit, V.E. Miron, J. Priller, S.J. Jenkins, G.E. Hardingham, M. Blurton-Jones, N.A. Mabbott, K.M. Summers, P. Hohenstein, D.A. Hume, and C. Pridans. 2019. Deletion of a Csf1r enhancer selectively impacts CSF1R expression and development of tissue macrophage populations. Nat Commun 10:3215.

Roy, D.S., Y. Zhang, M.M. Halassa, and G. Feng. 2022. Thalamic subnetworks as units of function. Nat. Neurosci. 25:140–153.

Rüb, U., K. Stratmann, H. Heinsen, D. Del Turco, E. Ghebremedhin, K. Seidel, W. den Dunnen, and H.W. Korf. 2016. Hierarchical Distribution of the Tau Cytoskeletal Pathology in the Thalamus of Alzheimer’s Disease Patients. Journal of Alzheimer’s disease: JAD 49:905–915.

Scheltens, P., B. De Strooper, M. Kivipelto, H. Holstege, G. Chetelat, C.E. Teunissen, J. Cummings, and W.M. van der Flier. 2021. Alzheimer’s disease. Lancet 397:1577–1590.

Serrano-Pozo, A., M.P. Frosch, E. Masliah, and B.T. Hyman. 2011. Neuropathological alterations in Alzheimer disease. Cold Spring Harb Perspect Med 1:a006189.

Shi, Y., K. Yamada, S.A. Liddelow, S.T. Smith, L. Zhao, W. Luo, R.M. Tsai, S. Spina, L.T. Grinberg, J.C. Rojas, G. Gallardo, K. Wang, J. Roh, G. Robinson, M.B. Finn, H. Jiang, P.M. Sullivan, C. Baufeld, M.W. Wood, C. Sutphen, L. McCue, C. Xiong, J.L. Del-Aguila, J.C. Morris, C. Cruchaga, I. Alzheimer’s Disease Neuroimaging, A.M. Fagan, B.L. Miller, A.L. Boxer, W.W. Seeley, O. Butovsky, B.A. Barres, S.M. Paul, and D.M. Holtzman. 2017. ApoE4 markedly exacerbates tau-mediated neurodegeneration in a mouse model of tauopathy. Nature 549:523–527.

Tetlow, A.M., B.M. Jackman, M.M. Alhadidy, P. Muskus, D.G. Morgan, and M.N. Gordon. 2023. Neural atrophy produced by AAV tau injections into hippocampus and anterior cortex of middle-aged mice. Neurobiol. Aging 124:39–50.

Tracy, T.E., and L. Gan. 2018. Tau-mediated synaptic and neuronal dysfunction in neurodegenerative disease. Curr. Opin. Neurobiol. 51:134–138.

Udeochu, J.C., S. Amin, Y. Huang, L. Fan, E.R.S. Torres, G.K. Carling, B. Liu, H. McGurran, G. Coronas-Samano, G. Kauwe, G.A. Mousa, M.Y. Wong, P. Ye, R.K. Nagiri, I. Lo, J. Holtzman, C. Corona, A. Yarahmady, M.T. Gill, R.M. Raju, S.A. Mok, S. Gong, W. Luo, M. Zhao, T.E. Tracy, R.R. Ratan, L.H. Tsai, S.C. Sinha, and L. Gan. 2023. Tau activation of microglial cGAS-IFN reduces MEF2C-mediated cognitive resilience. Nat. Neurosci. 26:737–750.

Wang, B., H. Martini-Stoica, C. Qi, T.-C. Lu, S. Wang, W. Xiong, Y. Qi, Y. Xu, M. Sardiello, and H. Li. 2024. TFEB–vacuolar ATPase signaling regulates lysosomal function and microglial activation in tauopathy. Nature neuroscience 27:48–62.

Wang, Y., and E. Mandelkow. 2016. Tau in physiology and pathology. Nat. Rev. Neurosci. 17:5–21.

Wesseling, H., W. Mair, M. Kumar, C.N. Schlaffner, S. Tang, P. Beerepoot, B. Fatou, A.J. Guise, L. Cheng, S. Takeda, J. Muntel, M.S. Rotunno, S. Dujardin, P. Davies, K.S. Kosik, B.L. Miller, S. Berretta, J.C. Hedreen, L.T. Grinberg, W.W. Seeley, B.T. Hyman, H. Steen, and J.A. Steen. 2020. Tau PTM Profiles Identify Patient Heterogeneity and Stages of Alzheimer’s Disease. Cell 183:1699–1713 e1613.

Wu, T., B. Dejanovic, V.D. Gandham, A. Gogineni, R. Edmonds, S. Schauer, K. Srinivasan, M.A. Huntley, Y. Wang, T.M. Wang, M. Hedehus, K.H. Barck, M. Stark, H. Ngu, O. Foreman, W.J. Meilandt, J. Elstrott, M.C. Chang, D.V. Hansen, R.A.D. Carano, M. Sheng, and J.E. Hanson. 2019. Complement C3 Is Activated in Human AD Brain and Is Required for Neurodegeneration in Mouse Models of Amyloidosis and Tauopathy. Cell Rep 28:2111–2123 e2116.

Yadav, N., A. Toader, and P. Rajasethupathy. 2024. Beyond hippocampus: Thalamic and prefrontal contributions to an evolving memory. Neuron 112:1045–1059.

Yamamori, H., S. Khatoon, I. Grundke-Iqbal, K. Blennow, M. Ewers, H. Hampel, and K. Iqbal. 2007. Tau in cerebrospinal fluid: a sensitive sandwich enzyme-linked immunosorbent assay using tyramide signal amplification. Neurosci. Lett. 418:186–189.

Yin, Y., D. Gao, Y. Wang, Z.H. Wang, X. Wang, J. Ye, D. Wu, L. Fang, G. Pi, Y. Yang, X.C. Wang, C. Lu, K. Ye, and J.Z. Wang. 2016. Tau accumulation induces synaptic impairment and memory deficit by calcineurin-mediated inactivation of nuclear CaMKIV/CREB signaling. Proc Natl Acad Sci U S A 113:E3773–3781.

Yoshiyama, Y., M. Higuchi, B. Zhang, S.M. Huang, N. Iwata, T.C. Saido, J. Maeda, T. Suhara, J.Q. Trojanowski, and V.M. Lee. 2007. Synapse loss and microglial activation precede tangles in a P301S tauopathy mouse model. Neuron 53:337–351.

Yu, G., L.-G. Wang, Y. Han, and Q.-Y. He. 2012. clusterProfiler: an R package for comparing biological themes among gene clusters. Omics: a journal of integrative biology 16:284–287.

Yushkevich, P.A., J. Piven, H.C. Hazlett, R.G. Smith, S. Ho, J.C. Gee, and G. Gerig. 2006. User-guided 3D active contour segmentation of anatomical structures: significantly improved efficiency and reliability. NeuroImage 31:1116–1128.

Zhang, Z., and Y. Zhao. 2022. Progress on the roles of MEF2C in neuropsychiatric diseases. Mol Brain 15:8.

